# Genetic and Epigenetic Determinants Establish a Continuum of Hsf1 Occupancy and Activity Across the Yeast Genome

**DOI:** 10.1101/306704

**Authors:** David Pincus, Jayamani Anandhakumar, Prathapan Thiru, Michael J. Guertin, Alexander M. Erkine, David S. Gross

## Abstract

Heat Shock Factor 1 (Hsf1) is the master transcriptional regulator of molecular chaperones and binds to the same *cis*-acting element - Heat Shock Element (HSE) - across the eukaryotic lineage. In budding yeast, Hsf1 drives transcription of ~20 genes essential to maintain proteostasis under basal conditions, yet its specific targets and extent of inducible binding during heat shock remain unclear. Here we combine Hsf1 ChIP-seq, nascent RNA-seq and Hsf1 nuclear depletion to quantify Hsf1 binding and transcription across the yeast genome. Hsf1 binds 74 loci during acute heat shock, 46 of which are linked to genes with strong Hsf1-dependent transcription. Most of these targets show detectable Hsf1 binding under basal conditions, but basal occupancy and heat shock-inducible binding both vary over two orders of magnitude. Notably, Hsf1’s induced DNA binding leads to a disproportionate (up to 50-fold) increase in nascent transcription. While variation in basal Hsf1 occupancy poorly correlates with the strength of the HSE, promoters with high basal Hsf1 occupancy have nucleosome-depleted regions due to the presence of ‘pioneer’ factors. Such accessible chromatin may be critical for Hsf1 occupancy of its genomic sites as the activator is incapable of binding HSEs embedded within a stable nucleosome *in vitro*. In response to heat shock, however, Hsf1 is able to gain access to nucleosomal sites and promotes chromatin remodeling with the RSC complex playing a key role. We propose that the interplay between nucleosome occupancy, HSE strength and active Hsf1 levels allows cells to precisely tune expression of the proteostasis network.

## Introduction

The cellular response to thermal stress is directed by a transcriptional program whose basic components - sequence-specific activator, *cis*-response DNA element and core target genes - have been conserved since the last common ancestor in the eukaryotic lineage (Wu 1995; Verghese et al. 2012). Heat Shock Factor 1 (Hsf1), master activator of the heat shock response, is a winged helix-turn-helix transcription factor that recognizes Heat Shock Elements (HSEs) located upstream of genes encoding chaperones, co-chaperones and other cytoprotective Heat Shock Proteins (HSPs).

HSPs maintain cellular protein homeostasis (proteostasis) and are required in higher concentrations under stressful conditions to deal with unfolded cytosolic and nucleoplasmic proteins. This is principally achieved via their Hsf1-mediated transcriptional up-regulation. In addition to its evolutionarily conserved role, mammalian HSF1 contributes to carcinogenesis through driving distinct transcriptional programs in both tumor cells as well as supporting stroma (Dai et al. 2007; Mendillo et al. 2012; Scherz-Shouval et al. 2014). HSF1 function has also been linked to normal development, neurodegenerative disease and aging (Neef et al. 2011; Li et al. 2017). Thus, gaining a deeper understanding of Hsf1 biology may inform development of novel approaches to modulate HSF1.

Hsf1 is subject to multiple layers of regulation, with two shared across multiple phyla. Under non-stressful conditions, Hsf1 exists primarily as a non-DNA-binding monomer, in either nucleus or cytoplasm. In response to thermal or other environmental stress, the protein trimerizes and acquires the capacity for high-affinity DNA binding. In higher eukaryotes, the Hsp90 and Hsp70 chaperones (and their co-chaperones) and the TRiC/CCT co-chaperone complex have been suggested as direct repressors of Hsf1 under basal conditions (Abravaya et al. 1992; Zou et al. 1998; Neef et al. 2014). According to this model, the chaperones are titrated away upon heat shock by unfolded or misfolded proteins, allowing Hsf1 to trimerize, bind its cognate HSEs and *trans*-activate *HSP* genes. Although certain details of Hsf1 regulation differ between metazoans and budding yeast (Sorger and Nelson 1989; Liu et al. 1997), it was recently shown that Hsp70 binds yeast Hsf1 and negatively regulates it, transiently releasing in response to thermal stress and then rebinding upon exposure to sustained stress, constituting a two-component feedback loop (Zheng et al. 2016; Krakowiak et al. 2018). Hsf1 has also been suggested to be the downstream effector of various signaling cascades. In the case of yeast, phosphorylation positively tunes Hsf1’s trans-activation of target genes independent of chaperone regulation (Zheng et al. 2016).

Hsf1-regulated, heat shock-responsive genes have served as a paradigm for understanding basic mechanisms of transcription. For example, the existence of paused RNA polymerase II (Pol II) at the 5’-end of genes was first identified at *Drosophila HSP70* (Rougvie and Lis 1988). Hsf1-regulated genes in *S. cerevisiae* use both SAGA and TFIID pathways for activation, although SAGA plays a dominant role (Ghosh and Pugh 2011; de Jonge et al. 2017; Vinayachandran et al. 2018). In addition, Mediator has been shown to be a key coactivator of Hsf1-driven transcription in both metazoans and yeast (Park et al. 2001; Fan et al. 2006; Kim and Gross 2013). Dynamic gene-wide eviction of nucleosomes and their subsequent re-deposition has been observed at *HSP* genes in both yeast and *Drosophila* (Zhao et al. 2005; Petesch and Lis 2008; Kremer and Gross 2009). Recently, Hsf1-regulated genes in yeast have been observed to undergo striking alteration in their local structure and engage in highly specific *cis-* and *trans*-intergenic interactions upon their transcriptional activation, coalescing into discrete intranuclear foci (Chowdhary et al. 2017) (Chowdhary et al, submitted).

In budding yeast, the Msn2 and Msn4 gene-specific transcription factors drive transcription of a large set of genes (200-300) in response to a variety of environmental stresses, including heat, oxidative, osmotic and salt stress (Gasch et al. 2000; Elfving et al. 2014). Msn2/Msn4-regulated genes include several *HSP* genes, but the Msn2/Msn4 regulon is by and large distinct from that controlled by Hsf1. Nonetheless, the identity of the genes whose acutely induced expression is under the direct control of Hsf1 remains unclear. Prior global localization studies of Hsf1 have either lacked resolution or did not evaluate Hsf1 occupancy beyond the basal or chronically induced state (Lee et al. 2002; Hahn et al. 2004; Solis et al. 2016; de Jonge et al. 2017).

Here we investigate the link between Hsf1 occupancy, its function and the epigenetic determinants underpinning its genome localization under basal, acute heat shock and chronic heat shock states in *S. cerevisiae*. Our results reveal that Hsf1 binds to a core set of 43 loci under control conditions; that the vast majority of these and 31 others are occupied at substantially higher levels following heat shock; and of these 74 bound loci, 46 are associated with genes whose transcriptional activation is Hsf1-dependent. Additionally, our analysis reveals a central role played by pre-set nucleosome positioning and the chromatin remodeler RSC in regulating the dynamic transcriptional response to heat shock in this organism.

## Results

### Yeast Hsf1 binds inducibly to the vast majority of its target genomic sites

To investigate Hsf1 DNA binding genome-wide, we performed ChIP-seq to determine where and how strongly it binds under basal (non-heat-shock (NHS)), acute heat shock (HS) and chronic HS states. To circumvent problems typically encountered when quantifying ChIP-seq data that combines dynamic protein binding with high coverage over a small genome, we performed parallel IPs using both anti-Hsf1 serum as well as pre-immune serum at all time points, and we generated paired-end sequencing data to determine the full sequence of the captured fragments (see Methods). Analytically, we subtracted signal from the matched pre-immune samples, used only properly paired reads and allowed for duplicate reads when quantifying fragment pileups. Such an approach revealed that Hsf1 binding could be detected under NHS conditions (30°C) and was centered ~200 bp upstream of the TSS. In cells exposed to acute heat shock (30° to 39°C shift for 5 min), its occupancy increased ≥4-fold genome-wide and remained elevated in cells chronically exposed (2 h) to the higher temperature (see metagene analysis in Figure 1A). Representative occupancy profiles are provided in Figure 1B, which illustrate both the specificity and reproducibility of our ChIP-seq analysis.

**Figure 1.**
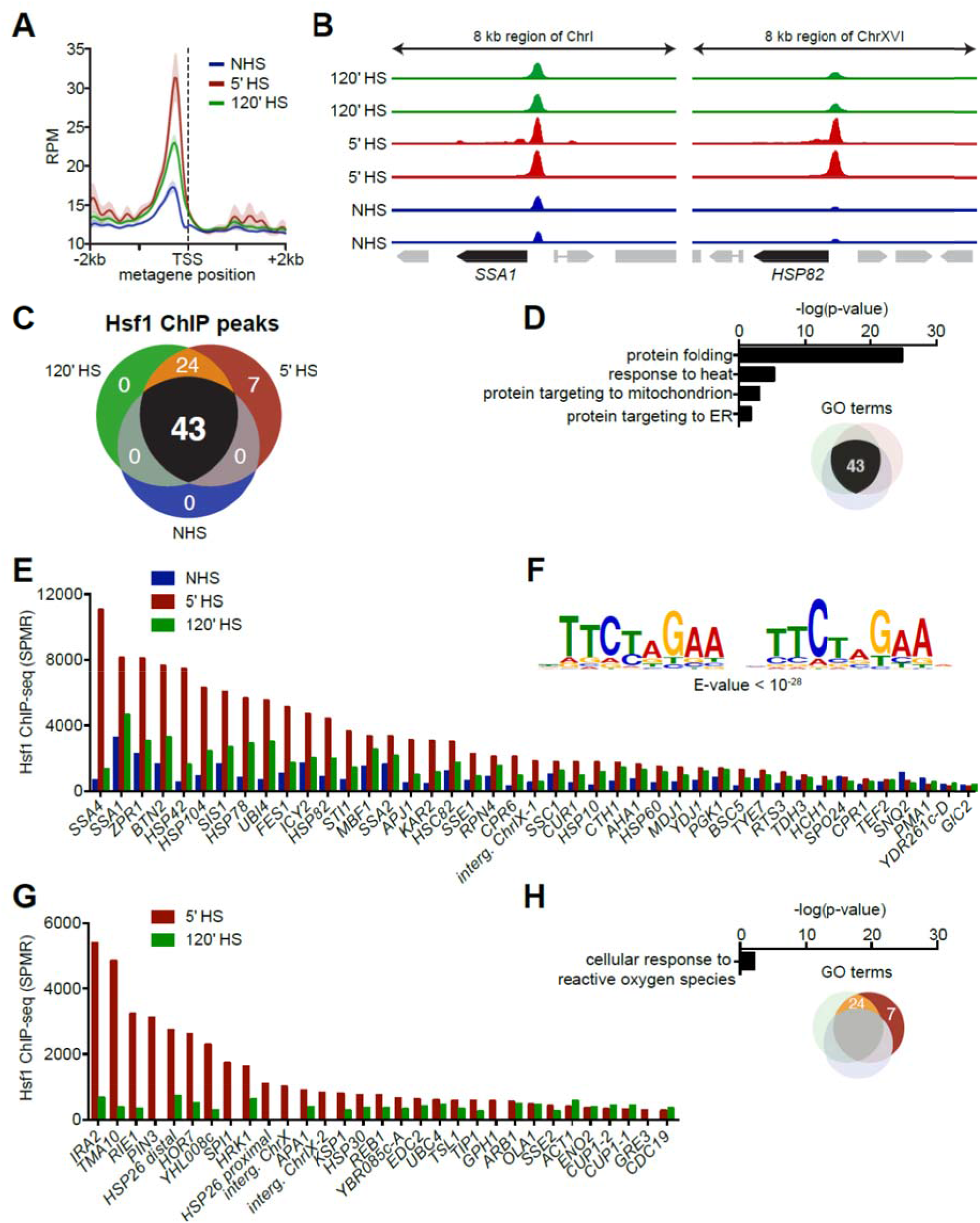
Hsf1 ChIP-seq reveals differential basal and heat shock-inducible binding across the Hsf1 regulon. **A)** Metagene plot of Hsf1 ChIP-seq signal genome-wide with respect to the transcription start site (TSS) under non-heat shock (NHS) conditions and following 5-min and 120-min heat shock (HS). Strain BY4741 was used for this and all other Hsf1 ChIP-seq assays in this work. RPM, reads per million mapped reads. **B)** Browser images of the *SSA1* and *HSP82* loci showing Hsf1 ChIP-seq signal in two biological replicates under NHS conditions and following 5-min and 120-min HS. The Y-axes are normalized to the maximum displayed signal in the 5-min time point. **C)** Venn diagram showing the number of Hsf1 ChIP peaks that surpassed the background cutoff in both biological replicates under each condition. **D)** Gene ontology (GO) term enrichment values for genes immediately downstream of the 43 ChIP peaks. **E)** Normalized Hsf1 ChIP signal at the 43 peaks identified under all three conditions. SPMR, signal per million mapped reads. **F)** Consensus motif identified under Hsf1 ChIP peaks detected under all conditions. **G)** Normalized Hsf1 ChIP signal at the 31 peaks detected only under HS conditions. **H)** GO term enrichment values for the 31 Hsf1 ChIP peaks detected only under HS conditions.

Two sets of Hsf1 genomic targets were identified. A core set, comprised of 43 loci, was occupied under all conditions, with occupancy typically increased in cells exposed to thermal stress (Figures 1C, 1E). A second set, comprised of 31 loci, showed subthreshold occupancy under NHS conditions but was inducibly occupied following acute HS; of these, 24 remained Hsf1-bound in cells chronically exposed to thermal stress (Figures 1C, 1G). We performed locus-specific ChIP to validate examples of each (Figure S1A). The core set of Hsf1 targets was enriched for proteostasis components in the cytosol, ER and mitochondria (Figure 1D). Notably, within these 43 core targets, we observed strong inducible binding with a broad range of inducible levels - up to 50-fold at select sites (Figure 1E). Indeed, all but five core targets exhibited elevated occupancy. Thus, yeast Hsf1 inducibly binds 69/74 of its genomic targets upon heat shock.

Analysis of the regions occupied by Hsf1 pulled out a sequence of 20 bp comprised of tandem inverted repeats of NTTCT as the most enriched motif (Figure 1F). This motif is consistent with previous characterization of HSEs from other organisms (Xiao et al. 1991; Leach et al. 2016; Vihervaara et al. 2017). Among the heat-shock-only Hsf1 binding sites, there was a modest enrichment for oxidative stress-responsive genes (Figure 1H) but no enrichment for an alternative binding motif (data not shown).

### DNA-bound Hsf1 is differentially active during basal and acute heat shock states

As previous genome-wide studies (Hahn et al. 2004; Eastmond and Nelson 2006; Solis et al. 2016; de Jonge et al. 2017) failed to define the Hsf1-dependent repertoire of target genes during acute heat shock, we were interested in knowing whether genes with strong heat shock-inducible and heat shock-only binding were in fact transcriptionally induced in an Hsf1-dependent manner. To test this, we deployed a yeast strain in which we could rapidly induce nuclear export of Hsf1 using the Anchor-Away system developed by Laemmli and colleagues (Haruki et al. 2008; Solis et al. 2016). Analysis of genome-wide transcription rates under NHS and 5 min HS states using nascent mRNA sequencing (NAC-seq; see Methods) revealed that the basal transcription of 18 genes was Hsf1-dependent, consistent with a previous study (Solis et al. 2016), while HS-induced transcription of this set of genes plus an additional 34 were also Hsf1-dependent (Figure 2A-D). Most Hsf1-dependent genes (46 / 52) were also occupied by Hsf1 in 5 min heat-shocked cells (Figure 2C), arguing that this group of genes, which is derived from both core and inducible categories (Figures 1E, G), is directly regulated by Hsf1. The remaining six exhibited sub-threshold Hsf1 occupancy. While not meeting our stringent cut-off (see Methods), these genes may be direct targets of Hsf1 as well (e.g., SSÆ3 [Figure S1A]). The targets identified by Hsf1 ChIP-seq whose NHS and 5 min HS transcription was Hsf1-independent are largely comprised of highly expressed housekeeping genes (Figure S2) that are presumably controlled by multiple gene-specific, functionally redundant activators. Notably, for the 46 genes corroborated by both ChIP-seq and NAC-seq, induced DNA binding by Hsf1 led to a disproportionate increase in nascent transcription during the first 5 min of HS (typically 7.5-fold and in certain cases >50-fold [Figure 2E]) (see Discussion).

**Figure 2.**
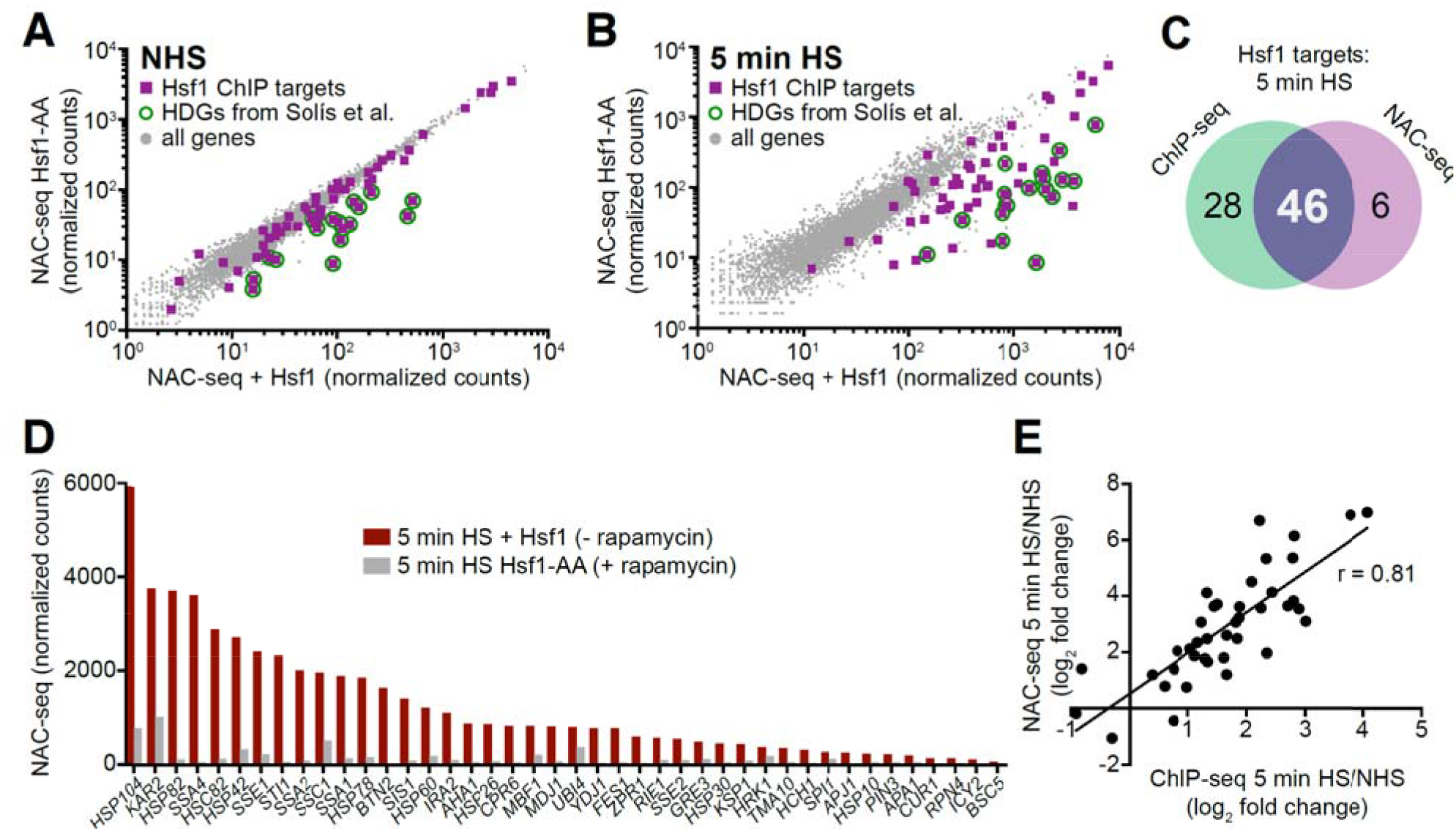
Nascent mRNA-seq (NAC-seq) coupled with Hsf1-Anchor Away reveals genes dependent on Hsf1 for their basal and induced transcription. **A)** NAC-seq counts transcriptome-wide under NHS conditions in the presence and absence of nuclear Hsf1 using the Hsf1 Anchor Away system (Hsf1-AA). 1 μM rapamycin was added for 45 minutes to deplete Hsf1 from the nucleus. Purple squares are genes with Hsf1 ChIP peaks; green circles show Hsf1 targets identified previously under NHS conditions (Solis et al. 2016). HDGs, Hsf1-dependent genes. **B)** As in (A), but following 5 min HS. **C)** Venn diagram comparing Hsf1 ChIP-seq targets and NAC-seq targets following a 5-min HS. **D)** NAC-seq counts for shared ChIP-seq/NAC-seq Hsf1 targets following a 5-min HS in the presence and absence of nuclear Hsf1. (Note: NAC-seq counts represent total nascent transcription and are not normalized for gene length (see Methods).) **E)** Scatter plot showing the correlation between HS-inducible Hsf1 DNA binding and HS-inducible transcription of shared ChIP-seq/NAC-seq Hsf1 targets.

### Hsf1 drives both unidirectional and bidirectional transcription at target loci

Examination of individual Hsf1-dependent genes revealed surprising variation with respect to their genomic arrangement and transcriptional response to DNA-bound Hsf1. Some, such as *HSC82* and *SSA2*, were exclusively activated by Hsf1 and, as revealed by Anchor-Away, entirely dependent upon it for their expression (Figure 3A; compare NAC-seq tracks-/+ Hsf1 in both NHS and 5 min HS samples). Others, such as genes within the bidirectional pairs *SIS1-LST8* and *YGR210C-ZPR1*, were symmetrically activated and also Hsf1-dependent (Figure 3B). On the other hand, the gene pairs *HSP82-YAR1* and *SSC1-TAH11* were asymmetrically activated, with one gene strongly induced while the other only weakly, yet both members were Hsf1-dependent (Figure 3C). In the case of *YAR1*, the Pol II transcript began well upstream of the gene, suggesting that this heat shock-inducible RNA might be non-coding. Since in each case the gene more strongly activated by Hsf1 was positioned closer to the Hsf1 site, the most parsimonious explanation is that proximity of the HSE to the core promoter/TSS dictates robustness of Hsf1 activation. The bidirectionally transcribed *MDJ1-HSP12* gene pair is striking for the strong Hsf1-dependence of one gene, *MDJ1,* but not of the other, *HSP12* (Figure 3D, Table S1). Indeed, *HSP12* appears to be primarily regulated by the alternative thermal stress-responsive activators Msn2 and Msn4 (Gasch et al. 2000). Also notable is the co-directionally oriented *NIS1-APJ1* gene pair. While both genes are heat shock-induced and Hsf1-dependent, *NIS1* is transcribed in the antisense direction (Figure 3E). Thus, Pol II transcripts arising from *YAR1* and *NIS1* are consistent with the idea that Hsf1 directs the expression of heat-shock-inducible lncRNAs.

**Figure 3.**
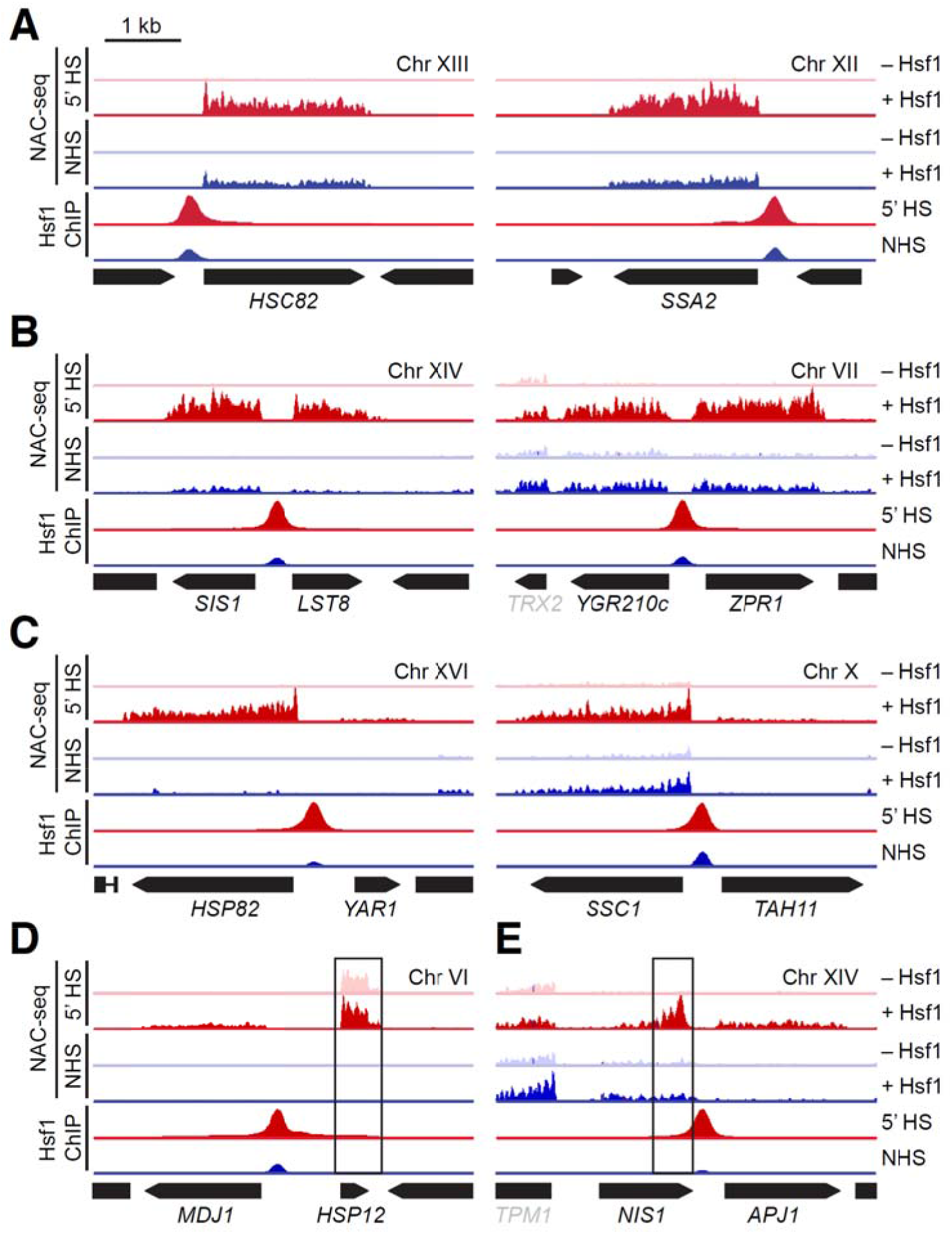
Hsf1 stimulates both unidirectional and bidirectional transcription. **A)** Browser images of 5-kb windows at Hsf1-dependent loci. Tracks show NAC-seq and Hsf1 ChIP-seq under NHS and 5-min HS conditions; NAC-seq was conducted in the presence and absence of nuclear Hsf1 using the Hsf1-AA system. ChIP-seq and NAC-seq tracks were normalized to the maximum displayed value for each locus in the 5-min heat shock sample. Hsf1 was anchored away with 1 μM rapamycin for 45 min. **B)** As in (A), but for loci that show near-stoichiometric, Hsf1-dependent bidirectional transcription. **C)** As in (B), but for loci that show sub-stoichiometric, Hsf1-dependent transcription of the non-chaperone gene. **D)** As in (C), but for the *MDJ1/HSP12* locus on chromosome VI. Although both *MDJ1* and *HSP12* are induced by heat shock, only *MDJ1* is Hsf1-dependent. **E)** As in (C), but for the *NIS1/APJ1* locus on chromosome XIV. There is HS-and Hsf1-dependent bidirectional transcription in the sense direction for *APJ1* and in the antisense direction for *NIS1*.

### A pre-set chromatin state poises Hsf1 for constitutive occupancy of core target loci

As we observed a broad spectrum of Hsf1 binding across the genome under both basal and heat-inducing conditions, we wished to know if one or more properties of the preset chromatin landscape correlated with this behavior. We examined these at three classes of Hsf1 target genes: (i) those with strong Hsf1 binding under NHS conditions (“High NHS Binding”); (ii) those with intermediate Hsf1 binding under NHS conditions (“Intermediate NHS Binding”); and (iii) those whose Hsf1 binding was only detectable following 5 min HS (“Low NHS Binding”).

High NHS Binding genes such as *SSA1, SSA2* and *HSC82* all displayed prominent nucleosome-free regions (NFRs) at the site of Hsf1 binding (Figure 4A; highlighted in yellow). In addition, nucleosomes flanking and downstream of these NFRs were enriched in histone marks linked to transcription, including acetylated H2A, H3, H4; H3 K4me3; H3 K36me3; and H3 K79me (Li et al. 2007). However, enrichment of these marks varied between genes. The pre-set chromatin landscape of Intermediate NHS Binding genes, epitomized by *HSP78, HSP82* and *HSP104*, was similar although the breadth of the NFR was noticeably reduced (Figure 4B). This may reflect the smaller proportion of cells with Hsf1 pre-bound upstream of these genes. Finally, the pre-set chromatin of Low NHS Binding genes, exemplified by *SSA4, HSP26* and *TMA10*, either lacked an NFR altogether (*SSA4*) or showed a greatly muted NFR (*TMA10, HSP26*) (Figure 4C). Instead, the upstream regions of these genes were enriched in histone acetylation marks, while classic methylation marks of transcription (H3K4me3, H3K36me36 and HK79me3) were depleted. Absence of the latter may reflect the fact that these genes express at very low levels in NHS cells as revealed by NAC-seq (Table S1). Notably, no class displayed enrichment of the Htz1 (H2A.Z) variant, contrary to current models which posit that this variant is enriched at nucleosomes flanking promoter-associated NFRs (Hartley and Madhani 2009).

**Figure 4.**
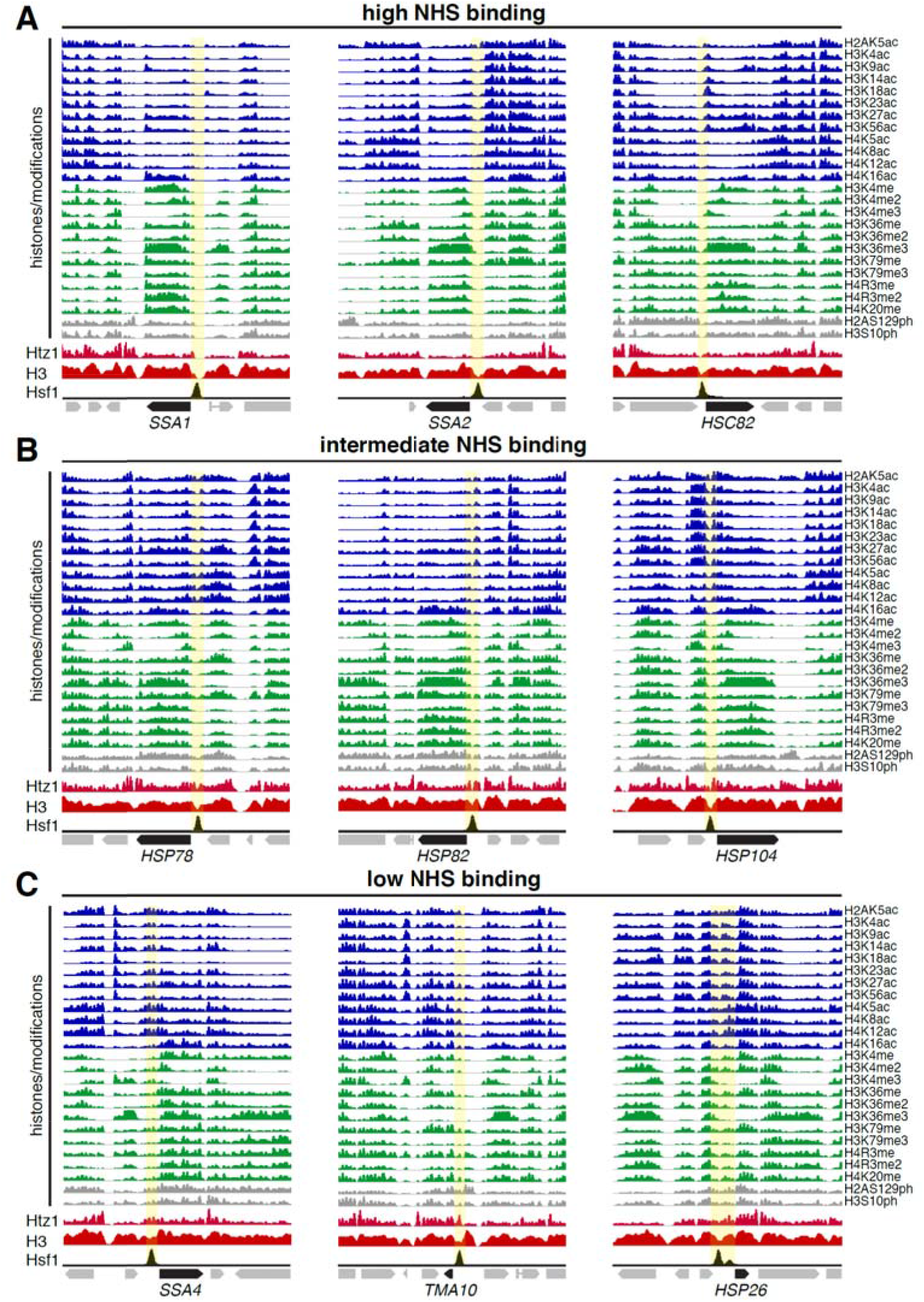
Hsf1’s constitutive occupancy correlates with pre-set accessible chromatin, while its inducible occupancy correlates with partial nucleosome occupancy and pre-existing histone acetylation. **A)** ChIP-seq profiles of histone acetylation, methylation and phosphorylation; H2A.Z (Htz1); total H3; and Hsf1 over 10 kb windows of the indicated High Hsf1 NHS Binding loci (high Hsf1 occupancy under control conditions). Location of NFR is highlighted in yellow. All tracks except Hsf1 were normalized to their own maximum displayed signal and are from publicly available NHS datasets. Hsf1, 5-min HS state (this study). See Methods for GEO accession numbers. **B)** As in (A), except depicted are Intermediate Hsf1 Binding loci (NHS state). Location of presumptive NFR in 5 min HS state is highlighted. **C)** As in (B), except Low Hsf1 Binding loci (NHS state).

### Hsf1 cannot bind HSEs embedded within a reconstituted, stably positioned nucleosome

The foregoing analysis suggests that a primary determinant of Hsf1 occupancy in NHS cells is whether or not the upstream region of a target gene is pre-assembled into stable chromatin. A powerful model to test this idea is the UAS region of the *HSP82* gene, which is occupied by Hsf1 under basal conditions (Figures 1E, S1A), while also partially assembled into chromatin, as inferred from histone ChIP and ChIP-seq assays (Figures 4B, 5A, and S7B). Upon acute HS, Hsf1 occupancy increases several-fold while nucleosome occupancy is reduced 80-90% (Figure S7) (Zhao et al. 2005; Kremer and Gross 2009).

**Figure 5.**
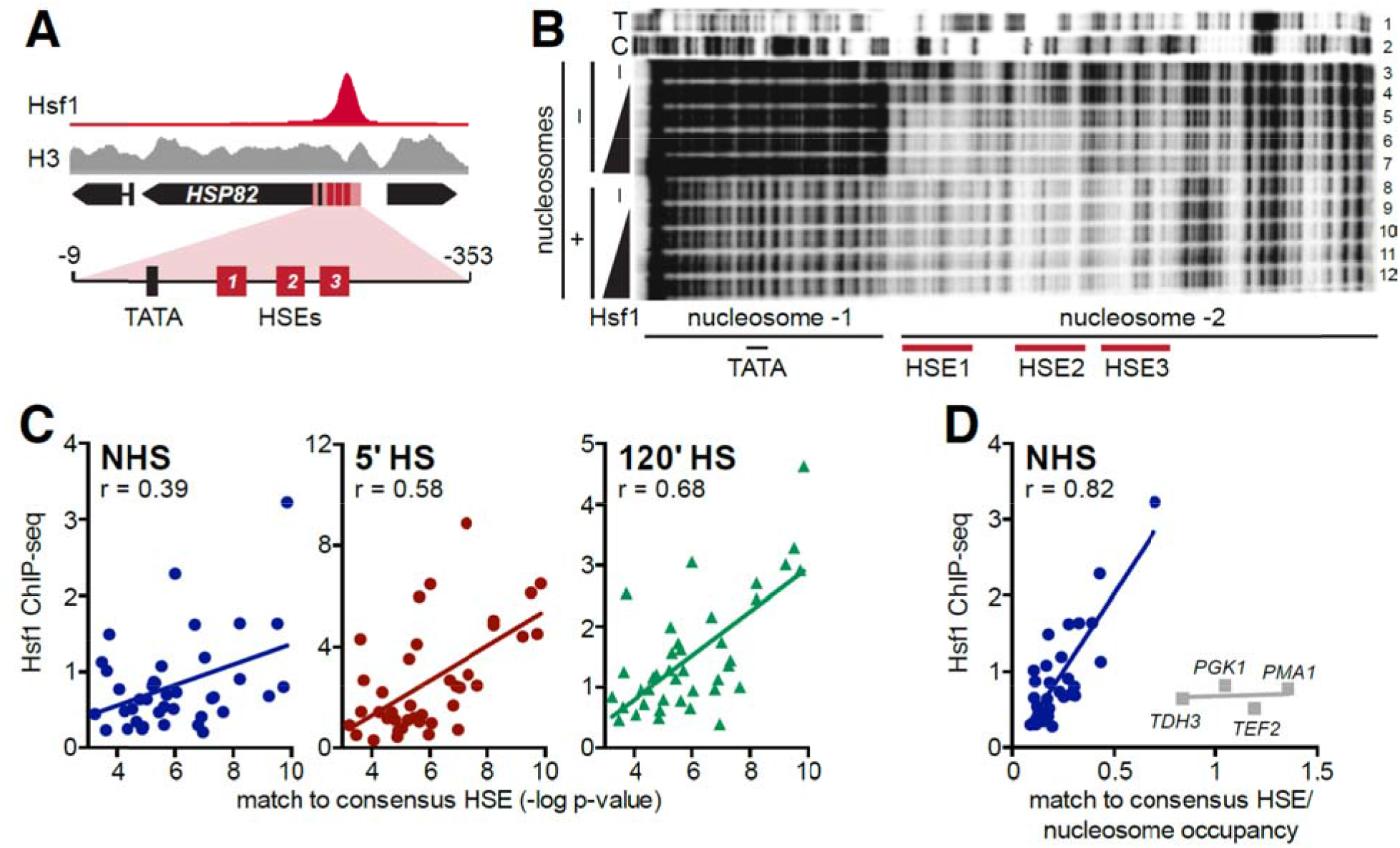
Hsf1 DNA binding is impeded by nucleosomes, both *in vitro* and *in vivo*. **A)** Hsf1 and H3 ChIP-seq signal at *HSP82* under NHS conditions. The region used for nucleosome reconstitution and DNase I footprinting is highlighted. **B)** DNA corresponding to the *HSP82* upstream region depicted in (A) (spanning- 9 to- 353 [ATG = +1]) and ^32^P-end labelled on the upper strand was either reacted directly with GST-Hsf1 (lanes 4-7) or following its reconstitution into a dinucleosome (lanes 912). Reconstitution was achieved using 1:1 (wt/wt) HeLa histone: DNA ratio and salt dilution, followed by purification over a glycerol gradient (see Figure S3). Both naked DNA and chromatin templates were challenged with increasing amounts of recombinant yeast Hsf1 (lanes 3 and 8 are-Hsf1 controls), then subjected to DNase I digestion. DNA was purified and electrophoresed on an 8% sequencing gel. **C)** Scatter plots of Hsf1 ChIP-seq signal as a function of the strength of the HSE for the NHS sample and the 5-and 120-min HS samples. HSE strength was determined by MEME as a p-value corresponding to how well the binding site beneath the summit of each ChIP peak matched the consensus HSE sequence (Figure 1F). **D)** As in (C), but here each p-value was divided by the H3 ChIP-seq signal below the summit of the Hsf1 peak under NHS conditions (H3 ChIP-seq data from Qiu et al. 2016). Outliers (gray) consist of Hsf1-independent housekeeping genes.

To directly test the ability of yeast Hsf1 to bind nucleosomal DNA, we reconstituted the upstream region of *HSP82* into chromatin using purified HeLa core histones (Figure S3, see Methods). Strikingly, this procedure results in the assembly of a dinucleosome, comprised of one nucleosome centered over the core promoter and a second over the UAS (HSEs1-3) and surrounding region (Figure 5B, lane 8). The nucleosome over the core promoter appears to be highly positioned (rotationally phased) as revealed by the chromatin-specific, ~10 base DNase I cleavage periodicity (compare lane 8 vs. lane 3) (Sollner-Webb et al. 1978). Equally striking, this footprinting pattern resembles that obtained by DNase I genomic footprinting of an inactivated allele of *HSP82* (*hsp82-ΔHSE1*) to which Hsf1 cannot bind and whose upstream region is occupied by two strongly positioned nucleosomes (Gross et al. 1993; Venturi et al. 2000). When we challenged this nucleosomal template with recombinant Hsf1, no Hsf1-dependent DNase I footprint could be detected (Figure 5B, lanes 9-12), in contrast to the naked DNA template where protection was evident over the entire UAS_HS_ (Figure 5B, lane 3 vs. 5-7).

Thus, although recombinant Hsf1 is capable of cooperative, high-affinity binding to the *HSP82* UAS (Figure 4B and (Erkine et al. 1999)), it cannot bind to the same DNA sequence when it is preassembled into a stably positioned nucleosome.

### Genome-wide Hsf1 occupancy is largely dictated by accessible, high-quality HSEs

While the above analysis suggests that pre-existing chromatin state plays an important role in dictating Hsf1 occupancy, other parameters, such as the quality of the HSE, may also play a role. To test this, we quantified how well the putative Hsf1-bound sequence under each ChIP peak matched the HSE consensus sequence we derived from all ChIP peaks (TTCTAGAAnnTTCTAGAA; Figure 1F) and compared this score (Figure S1B) with the amount of Hsf1 ChIP signal. We found little correlation under NHS conditions, but improved correlation as a function of time during heat shock (120' > 5' > NHS) (Figure 5C). Strikingly, when the quality of the HSE was considered in conjunction with nucleosome density as determined by total histone H3 ChIP-seq signal, a strong correlation was found between this (HSE quality/H3 signal) and Hsf1 occupancy under NHS conditions (Figure 5D, r = 0.81). That is, 64% of the variance in Hsf1 ChIP signal across the genome in the control state can be accounted for by this parameter (HSE quality/nucleosome density) alone.

### Pioneer transcription factors are enriched within upstream regions of genes with high NHS-bound Hsf1

How are nucleosome-free or nucleosome-depleted regions formed at Hsf1 binding sites? In mammals, "pioneer” transcription factors, epitomized by FoxA1 and related proteins, have been shown to potentiate the subsequent binding of gene-specific activators through their ability to invade repressive chromatin and create locally accessible regions (reviewed in Zaret and Carroll 2011; Zaret et al. 2016). The DNA binding proteins Rap1, Reb1 and Abf1 (sometimes termed General Regulatory Factors) have similar qualities: abundant, sequence-specific, constitutively DNA-bound and frequently located within nuclease-hypersensitive, open chromatin (Buchman et al. 1988; Chasman et al. 1990; Bai et al. 2011; Ganapathi et al. 2011). We therefore asked whether the genome-wide localization of Rap1, Reb1 and Abf1 in the NHS state showed any correspondence to that of Hsf1. As shown in Figure S4A, there is a striking enrichment of these factors overlapping or in close proximity to the nucleosome-depleted, high NHS binding Hsf1 sites of *SSA1* and *HSC82*. This relationship also exists with certain Intermediate NHS Binding Genes (e.g., *HSP82*) although not with others (e.g., *HSP104*) (Figure S4B). It is particularly evident at strongly expressed housekeeping genes such as *TEF2* and *TDH3* (Figure S4D); at such genes, Hsf1’s contribution to transcription is modest, especially under NHS conditions (Table S1). In marked contrast, the Low NHS Binding Genes *SSA4* and *HSP26* were located in Rap1/Reb1/Abf1 deserts with comparatively high H3 occupancy (Figure S4C). Together, this analysis suggests that pioneer factor binding is an important factor distinguishing high Hsf1 NHS binding targets from the low NHS binding, heat shock-inducible targets.

### Reb1 potentiates open chromatin and Hsf1-mediated transactivation and nucleosome displacement

To more directly address the role of pioneer factors in fostering Hsf1 binding and activity, we investigated the effect of mutating a high-affinity Reb1 binding site upstream of *HSC82*, creating a mutant allele termed *hsc82*-Δ*REB1*, and compared it to an isogenic mutant bearing a dual substitution of the two HSEs (termed *hsc82*-Δ*HSEs*) (Erkine et al. 1996). Hsf1 ChIP-seq and Reb1 ChEC-seq suggest high-level, overlapping occupancy of Reb1 and Hsf1 at *HSC82*^+^ (Figure 6A; (Zentner et al. 2015). Locus-specific ChIP confirms this, and moreover reveals that Reb1 occupancy is modestly diminished by a chromosomal substitution of HSE0 and HSE1 that obviates Hsf1 binding (Figures 6B and 6C). Similarly, chromosomal substitution of the Reb1 site, creating strongly reduced Reb1 occupancy (Figure 6B) and had a corresponding effect on Hsf1 occupancy in NHS cells (Figure 6C). Interestingly, Δ*REB1* strongly diminished *hsc82* expression, with transcript levels reduced >7-fold in control cells and 3-to 4-fold in acutely stressed ones (Figure 6D). The ability of Hsf1 to remodel *HSC82* chromatin was also impaired at *hsc82*-Δ*REB1*, as H3 occupancy was 3-4-fold higher throughout the gene in the mutant compared to WT at all time points. Notably, obviating Hsf1 binding led to an increase in H3 occupancy over both UAS and gene coding region.

**Figure 6.**
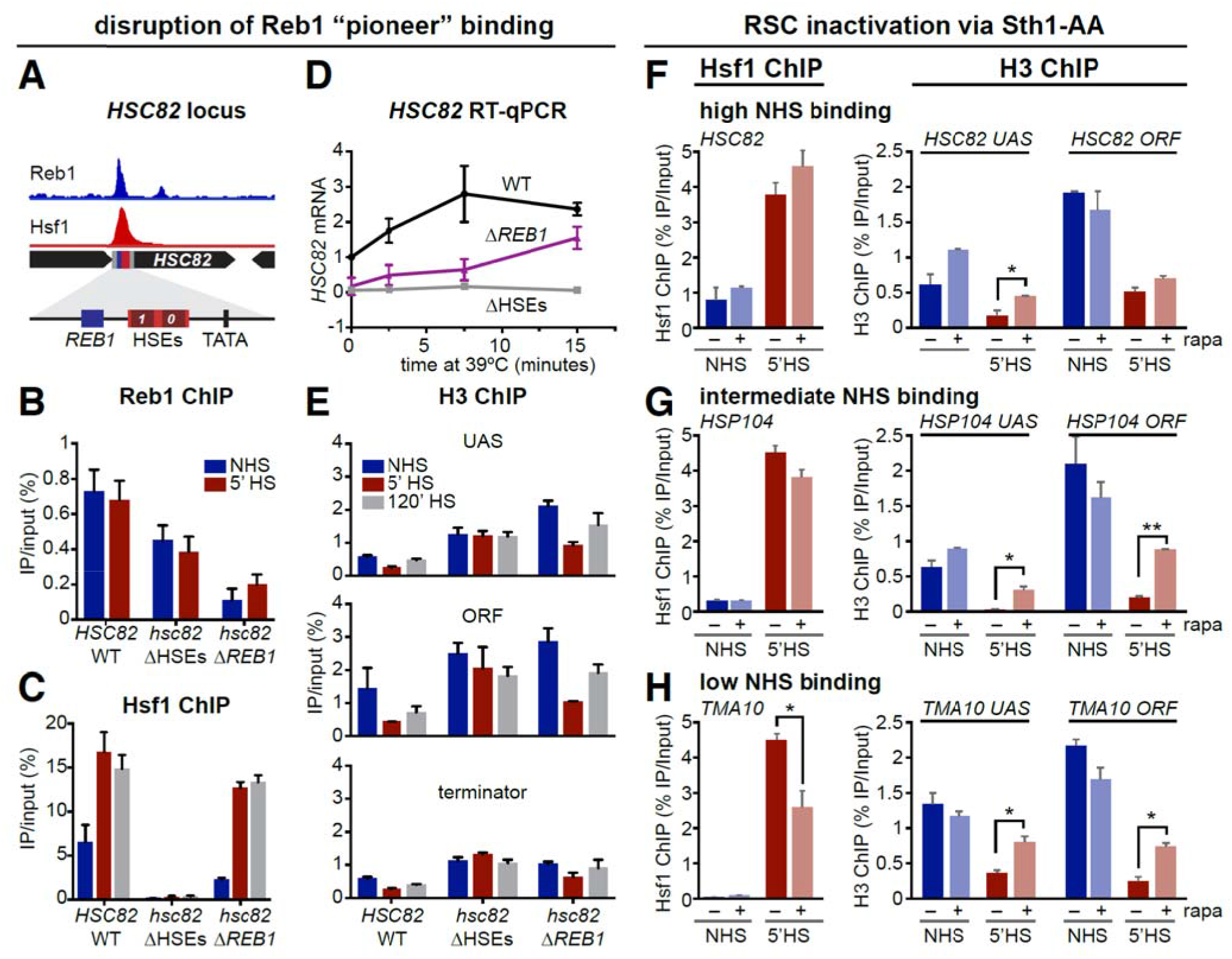
The pioneer factor Reb1 enables Hsf1 binding to a high NHS binding target while RSC cooperates with Hsf1 to displace nucleosomes during heat shock. **A)** Browser shot showing Reb1 ChEC-seq (Zentner et al. 2015) and Hsf1 ChIP-seq signal under NHS conditions at a 5 kb window around the *HSC82* locus. Both tracks are normalized to their maximum displayed values. Expanded view of the *HSC82* promoter shows the locations of the Reb1 binding site, HSEs and TATA box (centered at-249,-193 and -138, respectively; ATG = +1). **B)** Reb1-myc9 ChIP-qPCR analysis of the *hsc82* promoter under NHS and 5-min HS conditions in isogenic *HSC82, hsc82*-ΔHSEs and *hsc82*-Δ*REB1* cells conducted as described in Methods. The *hsc82*-Δ*HSEs* allele bears multiple point substitutions within HSE0 and HSE1; *hsc82*-Δ*REB1* bears a 10 bp chromosomal substitution of the Reb1 binding site (Erkine et al. 1996). Shown are means + SD (N=2 biological replicates; qPCR=4). **C)** Hsf1 ChIP analysis of the *hsc82* UAS region, conducted and analyzed as in (B). **D)** Quantification of *HSC82* mRNA expression level by RT-qPCR over a heat shock time course. Plotted are means +/- SD (N=2; qPCR=4). **E)** Histone H3 ChIP analysis of the *HSC82* promoter, mid-coding region (ORF) and 3’-UTR-terminator region. H3 ChIP signals were normalized to those detected at a nontranscribed locus, *ARS504*, which served as an internal recovery control. **F)** Hsf1 and H3 ChIP at *HSC82* under NHS and 5 min HS conditions, in the presence and absence of the RSC catalytic subunit, Sth1 (Sth1+ and Sth1-, respectively). Conditional nuclear depletion of Sth1 was achieved using an Sth1-AA strain that was pretreated with1 μM rapamycin for 2.5 h. Analysis and display as in **B** and **C**. **G)** As in (F), except *HSP104* was analyzed. **H)** As in (F), except *TMA10* was analyzed.

This by and large paralleled the effect of suppressing Reb1 (Figure 6E). Collectively, the data suggest that pioneer factors functionally cooperate with Hsf1 at select genes. In the absence of pioneer factor binding, other factors, including an increase in available Hsf1 (through chaperone titration as described above), are still able to drive nucleosome displacement upon acute heat shock (see model in Figure 7).

**Figure 7.**
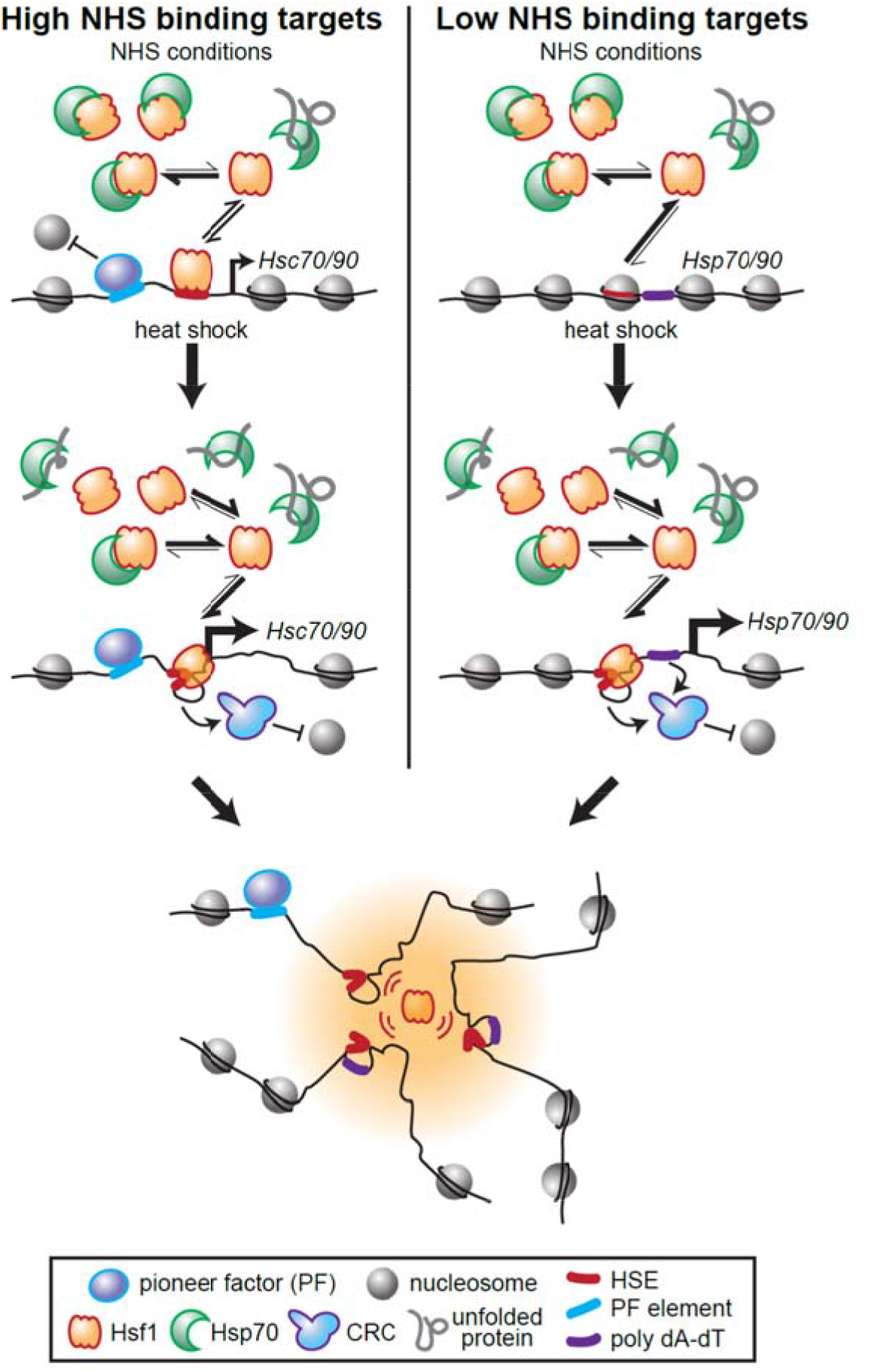
Model for differential basal and inducible Hsf1 binding across the genome. Target genes with high levels of Hsf1 binding under non-heat shock conditions (High NHS binding targets) have nucleosome-free (or depleted) regions upstream of the TSS due to the presence of pioneer factors. Such NFRs are occupied by the small fraction of DNA binding-competent Hsf1 present, stimulating basal transcription of linked *HSC70/90* genes (e.g., *SSA1, SSA2, HSC82*). These genes nonetheless show ~2-fold increases in Hsf1 binding upon acute heat shock when the bulk of Hsf1 is liberated from repressive association with Hsp70 (as unfolded proteins titrate Hsp70 away). This inducible binding further restructures the loci and drives increased transcription. In contrast, Low NHS binding targets (e.g., *SSA4, HSP26, TMA10*) lack proximal pioneer factor binding, and Hsf1 binding sites are occluded by nucleosomes. The fraction of free Hsf1 available under NHS conditions is insufficient to invade the chromatin at these sites. Upon heat shock, the large increase in DNA-binding competent Hsf1 allows Hsf1 to cooperatively bind its cognate HSEs, and along with poly(dA-dT) tracts that help recruit CRCs, to restructure the loci and drive high-level transcription. Active Hsf1 then drives looping and coalescence of its target loci into transcriptionally active foci that may form phase separated assemblies.

### RSC facilitates nucleosome depletion and Hsf1 binding within Rap1, Reb1, Abf1 deserts

Finally, we asked whether the abundant, ATP-dependent chromatin remodeling complex (CRC), Remodels Structure of Chromatin (RSC) (Cairns et al. 1996), plays a discernable role in creating a favorable chromatin template for Hsf1 binding and HS-dependent chromatin remodeling (Zhao et al. 2005; Zhang et al. 2014). RSC has previously been shown to play a critical role in remodeling a nucleosome covering Gal4 binding sites, rendering these sites accessible to Gal4 and as a consequence the linked *GAL1* and *GAL10* genes responsive to galactose (Floer et al. 2010). Using Anchor Away as above, we conditionally depleted Sth1, the catalytic subunit of RSC, from the nuclei of Sth1-FRB tagged cells, then subjected the cells to a subsequent acute heat shock (or not), and evaluated the effect on Hsf1 and H3 occupancy at representative High, Intermediate and Low NHS Binding Genes. Occupancy of either Hsf1 or H3 at the High NHS Binding gene *HSC82* was minimally affected by Sth1 perturbation (Figure 6F), whereas there was an increase in H3 over the UAS and coding regions of the Intermediate NHS gene *HSP104* in Sth1-depleted cells, although no effect on Hsf1 binding (Figure 6G). In contrast, both Hsf1 and H3 occupancy were affected by RSC depletion at the Low NHS Binding gene *TMA10* (Figure 6H). These results indicate that RSC plays an increasingly important, non-redundant role in Hsf1 regulation of genes whose pre-set chromatin structure is antagonistic to Hsf1 occupancy and function.

## Discussion

### Yeast Hsf1 inducibly binds DNA genome-wide in response to thermal stress

Results presented here indicate that *S. cerevisiae* Hsf1 binds heat-inducibly to the vast majority of its transcriptional targets (69/74). While seemingly contrary to original claims that yeast Hsf1 constitutively binds HSEs, based on EMSA (Sorger et al. 1987), genetic analysis (Jakobsen and Pelham 1988) or genomic footprinting (Gross et al. 1990), in fact our data validate that a small fraction of Hsf1 constitutively binds select HSEs even under control, NHS conditions (genome-wide occupancy ~20% of that seen following acute HS). This is consistent with the ability of yeast Hsf1 to trimerize under cell-free conditions (Sorger and Nelson 1989). Isolation of cell lysates or nuclei (prerequisite for EMSA and genomic footprinting) require spheroplasting, a procedure that induces the yeast heat shock response (Adams and Gross 1991), thereby providing a plausible explanation for the earlier claims.

Interestingly, Hsf1’s occupancy under NHS conditions, while correlating poorly with the quality of the bound HSE, shows a striking correlation to a related parameter, quality of HSE/nucleosome occupancy (r = 0.81) (compare Figures 5C and 5D). This is consistent with the fact that recombinant Hsf1 cannot bind HSEs assembled into a stable nucleosome (Figure 5B), even one that has been reconstituted with hyperacetylated histones (A.M.E. and D.S.G., unpublished observations). Thus, other factors must come into play under physiological conditions, and we demonstrate that a pre-set, nucleosome-free (or depleted) chromatin structure typifies the UAS/promoter regions of High NHS Binding Genes. One attribute underpinning the NFRs of high Hsf1 occupancy promoters is presence of constitutively bound "pioneer” factors such as Rap1, Reb1 and Abf1. These abundant DNA-binding proteins are overrepresented at the UAS/promoter regions of such genes, while unrepresented at Low NHS Binding Genes. They may work through recruitment of CRCs that mediate the nucleosome-free state (Krietenstein et al. 2016).

At Low NHS Binding Genes, relocalization of RSC that occurs upon heat shock (Vinayachandran et al. 2018) may be enhanced by the presence of poly(dA-dT) motifs (Lorch et al. 2014) that are significantly enriched within the NFRs of these genes (see Figure S5). Given that our evidence argues against a dominant role for RSC, Hsf1 likely recruits multiple CRCs (Zhao et al. 2005; Shivaswamy and Iyer 2008; Krietenstein et al. 2016). In addition, heat shock-activated Hsf1 triggers widespread changes in the genome that may contribute to its ability to occupy nucleosomal genomic sites and expand its regulon. Hsf1-target genes engage in frequent intergenic (both *cis-* and *trans-*) interactions during acute heat shock; such coalescence is strictly dependent on Hsf1 and encompasses High NHS, Intermediate NHS and Low NHS Genes (Chowdhary et al. 2017) (Chowdhary et al, in revision). *HSP* gene coalescence may be indicative of liquid-liquid phase separation postulated to underlie transcriptional control in higher eukaryotes (Hnisz et al. 2017). Thus, under this scenario, the high local concentration of Hsf1 liberated from Hsp70 upon heat shock would enable Hsf1 to gain a toe-hold at inducible targets and begin to recruit Mediator and other co-activators (many of which, like Hsf1 itself, harbor intrinsically disordered regions), driving coalescence among targets. These "Hsf1-bodies” could serve as a platform to recruit RSC and other CRCs through protein-protein interactions centered at coalesced UAS/promoters. A model illustrating the differential role played by pioneer factors and poly(dA-dT) tracts at High NHS Binding vs. Low NHS Binding Genes is presented in Figure 7.

### DNA-bound Hsf1 exists in multiple, functionally distinct states

An important implication of our study is that DNA-bound Hsf1 significantly differs in its ability to stimulate transcription of its targets depending on the proteotoxic state of the cell. Under control conditions, a core set of 43 genes is detectably occupied by Hsf1 (Figure 1E). Following acute HS, most of these 43 are occupied at substantially higher levels (genome wide Hsf1 occupancy increased ~4-fold). Nascent transcription of these genes increased even more, nearly 30-fold. Thus, DNA-bound Hsf1 exists in a far more active state in acutely stressed cells (~7.5-fold; Figure 5D). A possible basis for this is that the Hsp70 chaperone, shown to bind Hsf1 in whole cell extracts isolated from NHS cells but not from those exposed to 5 min HS (Zheng et al. 2016; Krakowiak et al. 2018), is also associated with DNA-bound Hsf1 under NHS conditions but not under conditions of acute heat shock (Figure 7). During more chronic exposures to stress, nascent transcription is reduced (Solis et al. 2016), and this may be accompanied by Hsp70 rebinding to DNA-bound Hsf1 (Zheng et al. 2016).

Related to the above, Hsf1-dependent transcription varies from one promoter to the next under a particular condition, even after normalization of ChIP-seq signal (Figure S6). What might account for this? Perhaps chromatin-associated factors, such as pioneer factors and/or chromatin remodeling complexes, either enhance or suppress the transcription-stimulating activity of DNA-bound Hsf1. A simple way this could be accomplished is by facilitating the association / disassociation of Hsp70. Especially intriguing is the hyperactive state of Hsf1 bound upstream of the essential, basal targets *SSA2* and *HSC82* (Figure S6A).

### Quality of HSE strongly correlates with Hsf1 binding in heat shock-induced cells

Earlier work on Hsf1 suggested that its binding avidity and transcriptional activity is linked to the presence of various types of HSEs, termed perfect, gap, step and direct repeat (reviewed in (Sakurai and Takemori 2007)). However, we have found that only when the role of chromatin is lessened - e.g., following nucleosome disassembly that accompanies heat shock (Zhao et al. 2005; Kremer and Gross 2009) - does a strong correlation exist between Hsf1 occupancy and quality of the HSE (Figure 5C). Indeed, 7/8 perfect HSEs located within promoter regions were occupied by Hsf1 upon 5 min HS (Table S2). While a far lower fraction of suboptimal HSEs were similarly occupied (e.g., 15/82 TTCnnGAAnnTTC motifs and 13/110 GAAnnTTCnnGAA motifs), there nonetheless exists a strong correlation between quality of the HSE and Hsf1 binding following acute heat shock, when chromatin remodeling is most prominent.

### A mutual antagonism exists between Hsf1 and nucleosomes under NHS conditions

A striking observation is that the *HSP82* upstream regulatory region can be reconstituted into two sequence-positioned nucleosomes using only core histones and DNA. This chromatin structure, which recapitulates the dinucleosome present within the upstream region of *hsp82-ΔHSE1* (Gross et al. 1993; Venturi et al. 2000) and resembles the dinucleosome reconstituted on a similar DNA sequence outfitted with p53 binding sites (Laptenko et al. 2011), may actively antagonize Hsf1 binding. Indeed, under either NHS or HS conditions, Hsf1 binding is not detectable at *hsp82-ΔHSE1*; instead, there is a 3-to 4-fold increase in histone H3 occupancy (Figure S7). At *HSP82*^+^ under NHS conditions, Hsf1 exists in a dynamic equilibrium with chromatin, as suggested by the cohabitation of Hsf1 with H2A, H2B, H3 and H4 (Figures 1E, 4B and S7) (Zhao et al. 2005), creating a chromatin structure hypersensitive to DNase I cleavage {Gross, 1990 #33}. Following acute heat shock, Hsf1’s cooperative binding of HSEs (Erkine et al. 1999), coupled with its recruitment of CRCs, HATs and other factors, culminates in domain-wide eviction of histones (Zhao et al. 2005; Kremer and Gross 2009; Kim and Gross 2013), and a 40-fold increase in nascent transcription (Table S1). Thus, the chromatin state of the *HSP82* promoter is subject to a dynamic remodeling process even under NHS conditions that is largely directed by Hsf1 itself.

### Relationship to previous genome-wide studies

It is instructive to compare findings reported here to previous genome-wide studies of Hsf1 occupancy. Using ChIP microarray analysis, Iyer, Thiele and colleagues reported that *S. cerevisiae* Hsf1 binds 210 genomic sites, 165 of which were located upstream of distinct open reading frames (ORFs) and the majority of which were heat-inducibly bound by Hsf1 (Hahn et al. 2004). Our ChIP-seq analysis, by contrast, identified only 74 genomic sites occupied by Hsf1, 69 of which were inducibly occupied and 46 of which were located upstream of genes whose heat-induced transcription was Hsf1-dependent. One explanation for the fewer genomic targets identified in our analysis is that we incorporated a high threshold for peak calls (see Methods). This led to the exclusion of six genes likely occupied by Hsf1 since their nascent transcription in acute heat-shocked cells is Hsf1-dependent and/or locus-specific ChIP detects above-background Hsf1 signal (Figure S1A). However, this fact alone cannot entirely explain the disparity between the datasets, since only 31/43 core Hsf1 targets and 14/31 inducible targets identified here were also identified in the earlier ChIP-chip analysis (Hahn et al. 2004). Holstege and colleagues examined Hsf1 occupancy in cells maintained under control, non-stressful conditions. Using ChIP-seq coupled with comparative dynamic transcription (cDTA) using Hsf1-AA strains, they identified 21 Hsf1 targets under NHS conditions (de Jonge et al. 2017); we identify these same 21 genes using a combination of ChIP-seq, NAC-seq and Anchor Away, plus four others (*SSA4, SSA2, HSP82* and *UBI4*).

Several previous studies examined the interplay of nucleosomes and Hsf1 binding. Similar to findings reported here, *Drosophila* HSF occupancy was found to be strongly linked to pre-set chromatin that is nucleosome-depleted, enriched in active histone marks, and whose landscape has pre-bound pioneer factors (Guertin and Lis 2010). Following its heat shock-induced binding, *Drosophila* HSF drives striking remodeling of chromatin (Petesch and Lis 2008), as we’ve observed here and elsewhere. In interesting contrast, Brown and colleagues reported little loss of histone over the UAS/promoter regions of Hsf1-bound loci in the pathogen *C. albicans* following exposure to heat shock (Leach et al. 2016). Why this is the case is unclear, but might be related to different abilities of the activation domains in *S. cerevisiae* Hsf1 vs. those in *C. albicans* to recruit CRCs.

Recently, Pugh and colleagues used ChIP-exo to map genome-wide protein-DNA interactions in *S. cerevisiae* under NHS and acute HS conditions at high resolution (Vinayachandran et al. 2018). They reported evidence for bidirectional transcription at Hsf1-regulated genes (based on the presence of symmetrically disposed PICs), as well as relocalization of RSC from coding regions to UAS/promoter regions in response to heat shock. Consistent with their findings, we observe bidirectional transcription at select Hsf1-bound loci in 5 min HS cells (Figure 3). We also find that RSC plays an important role in HS-dependent Hsf1 binding and histone eviction at Low NHS Binding genes (Figure 6H). As mentioned above, this correlates with an enrichment of poly(dA-dT) tracts within the NFRs of these genes (Figure S5). Our data are consistent with a model in which free Hsf1 cooperatively binds HSEs to recruit RSC and other CRCs, leading to the disassembly of nucleosomes in the UAS/promoter regions of Low NHS binding genes, and enabling high level Hsf1-dependent transcription during heat shock (Figure 7).

## Conclusions

The results we present here point to three clear conclusions: (1) Yeast Hsf1, like metazoan HSFs, inducibly binds to its target promoters across the genome during acute heat shock, contrary to popular models that presume the existence of a fundamental difference in DNA-binding behavior of yeast *vs.* metazoan HSF1. (2) There are a small number of essential chaperone genes to which Hsf1 binds strongly under NHS conditions; these genes have NFRs due to the presence of pioneer factors that expose strong HSEs. (3) Hsf1 expands its target regulon during heat shock by cooperating with CRCs to reveal HSEs occluded by nucleosomes, concomitant with driving its target genes - dispersed throughout the genome - into a coalesced, potentially phase-separated, state (Chowdhary et al. 2017). Together, these mechanisms allow cells to dynamically control the breadth and magnitude of the heat shock response to tune the proteostasis network according to need.

## Methods

### Yeast Strains

Strains AJ1001, AJ1002 and AJ1003 are derivatives of SLY101, HSE102 and GRF200 in which *REB1* was C-terminally tagged with the Myc9 epitope using integrative transformation as done previously (Kim and Gross 2013). Similarly, ACY101 was derived from HHY212 in which *STH1* was C-terminally tagged with the FRB domain (Haruki et al. 2008). All other strains have been previously described (see Table 1).

**Table 1.**
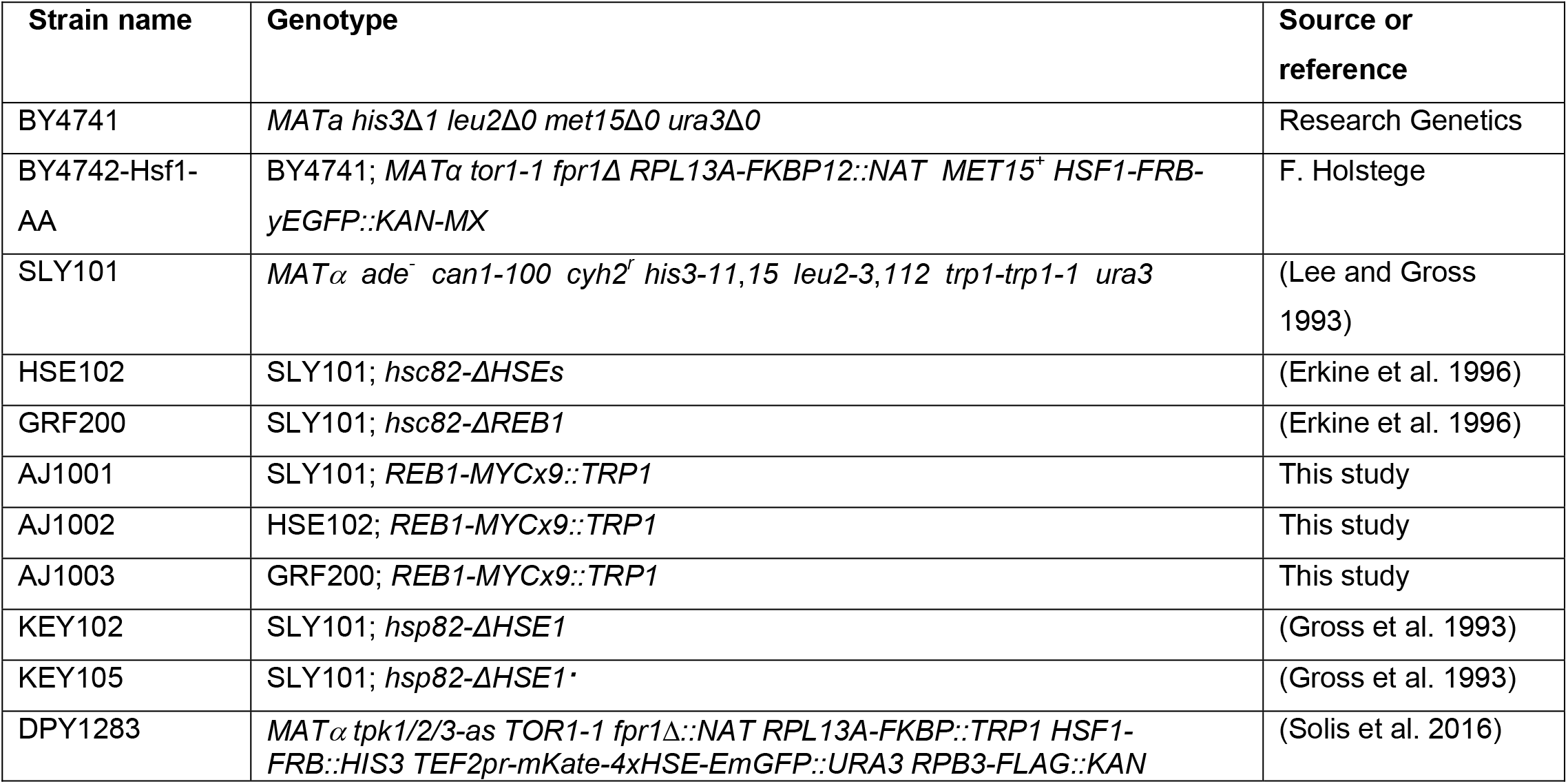
Yeast Strains.

### Chromatin Immunoprecipitation (ChIP)

ChIP experiments were performed as previously described (Kim and Gross 2013) except as noted below. 350-500 ml mid-log cell cultures were generally used in both rapamycin (final concentration 1 μg/ml) and heat shock time course experiments. 50 ml aliquots were removed at each time point to which formaldehyde was added to a final concentration of 1%. Cells were harvested, washed and resuspended in 250 μl lysis buffer, and lysed with vigorous shaking in the presence of glass beads (~300 mg) at 4°C for 30 min. Cell lysates were then transferred to 1.5 ml TPX tubes and sonicated at 4°C using a Diagenode Biorupter Plus (50 cycles with 30 sec pulses). This procedure generates chromatin fragments with a mean size of ~200-300 bp. TPX tubes were centrifuged to clarify supernatants which were then brought up to 2000 μl using ChIP lysis buffer. To perform immunoprecipitation, the equivalent of 500-800 μg chromatin protein (typically 200-400 μl) was incubated with one of the following antibodies: 1 μl of anti-Hsf1 (Erkine et al. 1996); 2.5 μl of anti-Myc (Santa Cruz Biotechnology); 1 μl of anti-H3 globular domain (Abcam; ab1791).

Immunoprecipitated DNA was resuspended in 60 μl TE; 2 μl were used in quantitative PCR (qPCR) with Power SYBR Green PCR Master Mix (Applied Biosystems 4367660) on an Applied Biosystems 7900HT Real-Time PCR system. DNA was quantified using a standard curve specific for each amplicon, and background signal arising from beads alone was subtracted. Background was determined by signal arising from incubating an equivalent volume of chromatin extract with Protein A Sepharose beads (GE Healthcare Life Sciences; CL-4B) and Protein G Sepharose beads (GE Healthcare Life Sciences; 4 Fast Flow). To normalize for variation in yield of chromatin extracts, in certain cases input was used. Briefly, 40 μl were removed from the 2000 μl chromatin lysate isolated as described above and the volume was brought up to 400 μl. HCHO-induced crosslinks were reversed and DNA deproteinized as for ChIP samples. Purified Input DNA was dissolved in 300 μl TE, and 2 μl were removed for qPCR. The signal arising from this represents 2% of total input chromatin. In the case of H3 ChIP, signal at a test locus was normalized to that obtained at ARS504 as described (Zhang et al. 2014). Primers used for detection and quantification of genomic loci in ChIP and Input DNAs are listed in Table S3.

### ChIP Sequencing (ChIP-Seq)

ChIP-Seq experiments were performed as above except as noted below. 600 ml mid-log cell cultures were used for each heat shock time course sample (0 min, 5 min and 120 min HS), formaldehyde was added to a final concentration of 1%. Cells were harvested, washed and 50 ml aliquots resuspended in 250 μl lysis buffer, and lysed with vigorous shaking in the presence of glass beads (~300 mg) at 4°C for 30 min. Cell lysates were then transferred to 1.5 ml TPX tubes and sonicated at 4°C using a Diagenode Biorupter Plus (60 cycles with 30 sec pulses). This procedure generates chromatin fragments with a mean size of ~100 - 250 bp. TPX tubes were centrifuged to clarify supernatants, which were then brought up to 3000 μl using ChIP lysis buffer. To perform immunoprecipitation, 100 μl was incubated with 4 μl of anti-Hsf1 antiserum (Erkine et al. 1996) and additionally, as control, 4 μl pre-immune serum used for each of the above three time points (derived from the same rabbit as the anti-Hsf1 immune serum). Five separate IPs were conducted for both the Hsf1 ChIP and the pre-Immune ChIP using 40 μl Protein A-Sepharose beads for each. Following purification of the IP and deproteinization, each ChIP DNA was suspended in 60 μl TE; the five samples were combined into one 300 μl pooled sample and quantified using Qubit Fluorimeter assay. 5 ng ChIP DNA were used to generate barcoded libraries using NEBNext Ultra DNA Library Prep Kit for Illumina (NEB E7370). Libraries were sequenced on lllumina Mi-Seq and NextSeq 500 instruments located in the LSUHSC Core Research Facility. Raw data are temporarily deposited on Dropbox but will be submitted to GEO prior to b publication. See links below (Data Access).

### Hsf1 ChIP-seq Analysis

Reads were aligned to the SacCer3 build of the yeast genome using Bowtie2. We used MACS2 to call and quantify peaks, filtering for properly paired reads and subtracting the pre-immune serum, and to generate a signal file that outputs fragment pileups per million reads in bedgraph format (signal per million mapped reads; SPMR).

It was important that we allowed MACS2 to model the expected value of duplicate reads (-keep dup auto) due the compact size of the genome and the high coverage of the dataset. Only peaks that had SPMR summit values above the background threshold of 250 were considered. Motif enrichment analysis was performed using MEME-ChIP. The MACS command used for a generic sample/pre-immune pair that had been fragmented to 180 bp:

> macs2 callpeak -t SAMPLE.bam -c PREIMMUNE.bam -f BAMPE -g 1.2e7 -n SAMPLE -p 1e-3 -nomodel -B -SPMR -extsize 180 -keep-dup auto

### Analysis of Previously Published Datasets

Histone modification and Htz1 ChIP-seq data was obtained from GSE61888 (Weiner et al. 2015). Histone H3 data are from GSE74787 (Qiu et al. 2016). Pioneer factor ChEC-seq data are from GSE67453 (Zentner et al. 2015). Data were visualized in the Integrated Genome Viewer (IGV).

### Nascent Transcript Sequencing (NAC-seq)

Nascent mRNA was enriched by purifying RNA that co-precipitates with RNA Pol II-component Rpb3. Hsf1 Anchor-Away cells expressing Rpb3-FLAG were grown at 30°C to 0D600 = 0.8 in YPD, treated +/- 1 μM rapamcyin for 45 minutes prior to harvesting before or after a 5-minute heat shock at 39°C. Cells were collected and lysed in a coffee grinder as described (Zheng et al. 2016) and resuspended in 1 ml IP buffer (20 mM HEPES pH 7.4, 110 mM KOAc, 0.5% Triton X-100, 0.1% Tween-20, 10 mM MnCl2, 1x Complete mini EDTA-free protease inhibitor, 50 units/ml SuperaseIn). DNA was digested with 150 units/ml RNAse-free DNAse for 20 minutes on ice and samples were centrifuged at 20,000 x g. Clarified lysate was added to 50 μl pre-washed anti-FLAG M2 magnetic beads and incubated for 2 hours at 4°C. Beads were washed 4x with 1 ml IP buffer and eluted in 2 x 50μl of 3xFLAG peptide (0.5 mg/ml) in IP buffer. RNA was purified using the Qiagen miRNeasy kit and libraries were prepared using the NEB Next Ultra RNA kit. Illumina sequencing libraries were sequenced at the Whitehead Genome Technology Core. Reads were aligned using Tophat, quantified with HTSeq-Count and normalized using DESeq2. Raw data are deposited at GEO: See link at Data Access.

### Reverse Transcription - Quantitative PCR (RT-qPCR)

Cells were cultivated in 600-650 ml mid-log cultures, and 50 ml aliquots were removed for each heat shock time point and treated with 1/100^th^ volume of 2 M sodium azide to terminate transcription. Total RNA extracted using phenol-chloroform and purified using an RNeasy kit (Qiagen 74204). 0.5 μg to 2 μg of purified RNA and random primers were used in each cDNA synthesis using the High-Capacity cDNA Reverse Transcription kit (Applied Biosystems 4368814). Synthesized cDNA was diluted 1:20 and 5 μl of diluted cDNA were added to each 20 μl real-time PCR reaction. Relative cDNA levels were quantified by the delta delta (ΔΔ) Ct method. The Pol III transcript *SCR1* was used as a normalization control for quantification of *HSP* mRNA levels. PCR primers used to detect cDNAs are provided in Table S3.

### Nucleosome Reconstitution and Hsf1 *In Vitro* Binding Assay

For nucleosomal reconstitution of the *HSP82* upstream regulatory region, we amplified a 344 bp region spanning -353 to -9 (where +1 = ATG start codon) by PCR using a plasmid (KEM101) harboring the *HSP82 Eco* RI fragment (-1359 to +1543) as template. The forward primer was end-labeled and gel-purified. Reconstitution was achieved using the salt dilution method. Briefly, a 20 μl reaction containing 5 μg (2 x 10^6^ cpm) of PCR fragment, 5 μg of unlabeled *Hae* III/*Msp* I-digested lambda DNA, and 10 μg HeLa histones were incubated in 15 mM Tris-HCl, pH 7.5, 5 M NaCl, 0.2 mM EDTA, 0.2 mM PMSF for 20 min at 37°C, then 5 min at RT. At 10 min intervals thereafter, 10 μl of 15 mM Tris-HCl, pH 7.5, 0.2 mM EDTA and 0.2 mM PMSF were added to a final volume of 200 μl. This mixture was then loaded onto a 7.5%-35% glycerol gradient (3.75 ml total volume) and spun at 33,500 rpm in a Beckman L8 rotor for 15 h at 4°C. The gradient was fractionated into 25 equal volumes and 5 μl from each fraction were electrophoresed on a 5% polyacrylamide TBE gel. The fraction containing pure dinucleosomes (no free DNA, no chromatin aggregates) was used for footprinting.

To conduct DNase I footprinting, we challenged the template, either free DNA or purified dinucleosome, with recombinant Hsf1. GST-Hsf1, purified from *E. coli* as previously described (Erkine et al. 1999), was pre-incubated with ~125 ng of DNA (50,000 cpm) of dinucleosomal template in 15 mM Tris-HCl, pH 7.5, 50 mM KCl, 4 mM MgCl_2_ for 20 min at RT (0, 0.5, 1.5, 4.5, 13.5 ng GST-Hsf1 were used; see Figure 5B). In parallel, ~125 ng of naked DNA were similarly incubated with GST-Hsf1. 5 μl of a 1/100 dilution of DNase I (Sigma D7291; 100 U/ μl stock) were then added to the mixture and digestion was allowed to proceed for 2 min. It was terminated by addition of EDTA to 5 mM.

## Data Access

ChIP-seq and NAC-seq datasets can be accessed at:

https://www.dropbox.com/sh/588d94asnn3ww6e/AADIgsf5RrhqSRc7m-bIUrgsa?dl=0

https://www.dropbox.com/sh/0e7ca339kzterqk/AADxScIlK0Byls-VBpEYzBDRa?dl=0

## Acknowledgements

We thank Amol Kainth for critical reading of the manuscript; Paula Polk (LSUHSC Core Research Facility) for assistance with ChIP-seq; Tom Volkert and the Whitehead Institute Genome Technology Core for assistance with Nascent Transcript sequencing (NAC-seq); Apeng Chen for strain construction; Jodi Taylor for assistance with data analysis; and Frank Holstege for yeast strains. This work was supported by an NIH Early Independence Award (DP5 OD017941) to D.P., a Feist-Weiller Postdoctoral Grant awarded to J.A., and grants from the NSF (MCB-1518345) and NIH (GM 128065) awarded to D.S.G.

## Author Contributions

D.P., J.A. and D.S.G. designed the project. J.A., D.P. and A.M.E. conducted the experiments. D.P., P.T. and M.G. performed the bioinformatics analysis. D.P., J.A., A.M.E. and D.S.G analyzed the data. D.P., J.A. and A.M.E. made the figures. D.S.G. and D.P. wrote the paper.

## Supplemental Figures

**Figure S1. Locus-specific analysis of Hsf1 binding and genome-wide ranking of its binding sites.**

**A)** Hsf1 ChIP analysis of the *HSP82* and SSA3 promoters over a heat shock time course in the presence or absence of nuclear Hsf1 using strain BY4742-Hsf1-AA. 1 μM rapamycin was added for 45 min to deplete Hsf1 from the nucleus prior to heat shock. Error bars represent SD of two biological replicates that are each comprised of two technical replicates. While SSA3 did not pass the threshold for Hsf1 binding by ChIP-seq, this analysis demonstrates a modest but significant induction of Hsf1 binding.

**B)** Hsf1 target binding sites ranked by their match to the consensus HSE sequence. P-values were computed by MEME. Demonstrates that there is no direct relationship between the strength of the HSE and the amount of Hsf1 bound. For example, the HSE of *SSA4*, which has very low Hsf1 occupancy under NHS conditions, is higher-ranking than that of SSA2, which has relatively high Hsf1 occupancy under the same conditions. Stronger correlations between HSE strength and Hsf1 occupancy exist for acute and chronic HS states (see Figure 5C).

**Figure S2. Housekeeping genes are occupied by Hsf1 but their expression is Hsf1-independent.**

Browser tracks show Hsf1 ChIP-seq under NHS and 5-min HS conditions, and NAC-seq in the presence and absence of nuclear Hsf1 (Hsf1-AA system) under NHS and 5-min HS conditions. ChIP-seq and NAC-seq tracks are normalized to the maximum displayed value in the 5-min HS sample. Hsf1 was anchored away with 1 μM rapamycin for 45 min.

**Figure S3. Reconstitution of the *HSP82* upstream region into chromatin.**

**A)** Purified HeLa histones were resolved on a 15% polyacrylamide gel containing 0.9 M acetic acid, 6 M urea and 0.37% Triton X-100 and stained with Coomassie blue.

**B)** 7.5 - 35% glycerol gradient analysis of reconstituted nucleosomes at the indicated ratios of core histones: DNA (wt/wt). DNA template, comprised of the *HSP82* upstream region spanning -353 to -9, was ^32^P-end-labeled on its upper strand.

**Figure S4. Pioneer transcription factor binding is a hallmark of promoters with high NHS Hsf1 occupancy.**

**A)** Browser images of 5 kb regions around representative genes with high Hsf1 occupancy under NHS conditions. Hsf1 ChIP-seq tracks, displayed for both NHS and 5 min HS conditions, are normalized to the maximum displayed value in the HS sample. The histone H3 ChIP-seq track is normalized to its maximum displayed value. The Abf1, Rap1 and Reb1 ChEC-seq tracks are plotted on an absolute scale from 0-15 reads per million mapped reads (RPM) (data from (Zentner et al. 2015)).

**B)** As in (A), except profiles of intermediate Hsf1 occupancy genes are displayed.

**C)** As in (A), except profiles of low Hsf1 occupancy genes are displayed.

**D)** As in (A), except displayed are profiles of representative Hsf1-independent housekeeping genes.

**Figure S5. Poly(dA-dT) tracts are significantly enriched near Hsf1 binding peaks at Low NHS Binding Genes.**

Consensus motif identified by MEME-ChIP using the 31 targets that displayed heat shock-only Hsf1 binding (see Figure 1G). Motif shown was identified within a window of 250 bp centered at the Hsf1 peak. In addition to these heat shock-only targets, Intermediate NHS Binding targets such as *HSP104, HSP82* and *HSP42* also harbor poly(dA-dT) tracts near their HSEs.

**Figure S6. Hsf1 binding weakly correlates with Hsf1-dependent transcription.**

**A)** Absolute Hsf1-dependent transcription under NHS conditions, defined as the normalized NAC-seq counts for a given gene in control cells minus the NAC-seq counts in cells with Hsf1 anchored away, plotted as a function of Hsf1 ChIP-seq signal at the promoter. Under these conditions, *SSA2* and *HSC82* show much greater Hsf1-dependent transcription than other genes.

**B)** As in (A), except 5 min heat shock state.

**Figure S7. Occupancy of Hsf1 at** ***HSP82*** **is obviated by chromosomal deletion of HSE1, with concomitant assembly of the UAS region into stable chromatin.**

**A)** Hsf1 ChIP analysis of the *hsp82* UAS region in isogenic *HSP82*^+^, *hsp82-ΔHSE1* and *hsp82-ΔHSE1*· strains (SLY101, KEY102 and KEY105, respectively) under 0, 5, 120 min heat shock conditions. The *hsp82-ΔHSE1* allele bears a 32 bp chromosomal deletion spanning HSE1 and flanking sequence; *hsp82-ΔHSE1*· bears a 32 bp substitution of the same region (Gross et al. 1993).

**B)** As in (A), except histone H3 density of the *hsp82* UAS region was evaluated. H3 ChIP signal was normalized to that detected at *ARS504* as described above (see Methods). Differential histone occupancy of *HSP82*^+^, *hsp82-ΔHSE1* and *hsp82-ΔHSE1*· suggest that ~70% of *HSP82*^+^ cells within the population are occupied by Hsf1, rather than by nucleosomes, over the UAS/promoter region.

